# Distinct genetic requirements underlie LITE-1-dependent photosensation and avoidance of high concentration of diacetyl in *C. elegans*

**DOI:** 10.64898/2026.10.13.758841

**Authors:** Alan Koh, Changchun Chen, André EX Brown

## Abstract

The eyeless nematode *C. elegans* can detect and respond to aversive blue and ultraviolet light using the gustatory receptor homolog LITE-1. In addition to its role in phototransduction, LITE-1 also functions as a chemoreceptor for the odorants diacetyl and 2,3-pentanedione. However, whether these different sensory modalities rely on the same downstream signalling components is unknown. In this study, we provide behavioural evidence that unlike its LITE-1-dependent photosensory function which relies on cGMP signalling components, its chemosensory function remains intact in mutants with disrupted cGMP signalling.

## Introduction

The ability to detect and respond to environmental stimuli is fundamental across all domains of life. Organisms have evolved a wide array of receptors to sense and interpret diverse external cues. Despite possessing a compact nervous system comprising only 302 neurons, the nematode *Caenorhabditis elegans* has evolved an impressive repertoire of sensory receptors [1, 2]. Among these are olfactory receptors that enable the detection of a broad range of volatile organic compounds, triggering behavioural responses that are often concentration dependent [1, 3, 4]. One such example is the compound diacetyl which at low concentration is attractive to *C. elegans* and is sensed via the olfactory receptor ODR-10 [5-7]. In contrast, at high diacetyl concentration, the signal becomes aversive, and an avoidance behaviour is mediated mainly by LITE-1 via chemosensory neurons ASK, ADL and possibly ASH via SRI-14 [8, 9].

LITE-1 is best known as a photoreceptor mediating light avoidance in *C. elegans* that belongs to the gustatory receptor family [10]. It was originally identified through genetic screens for mutants defective in light avoidance behaviour [11]. LITE-1 is sequentially and structurally different from known light sensing proteins such as opsins, including rhodopsin [11-13]. LITE-1 adopts an inverted membrane topology, differing from classical G protein-coupled receptors (GPCRs), highlighting its noncanonical nature [12]. Recent findings suggest that LITE-1 may function as a light-gated ion channel, activated by ultraviolet and blue light [11, 12, 14, 15]. Furthermore, LITE-1 is also required for the response to X-ray stimulation, although the underlying mechanism is unknown [16].

LITE-1 acts in several light-responsive sensory neurons including ASK, ASH, ADL, ASJ, and the interneuron AVG [13, 15, 17-19]. Signal transduction in ASJ neurons involves a well-characterised cGMP signalling cascade requiring the G proteins GPA-3 and GOA-1, guanylate cyclases ODR-1 and DAF-11, phosphodiesterases PDE-1,2,3 and 5, and signalling through the second messenger cGMP [13, 15]. The increase in cGMP levels activates cyclic nucleotide-gated (CNG) channels TAX-2 and TAX-4, driving an avoidance response (Fig. 1A) [13, 15]. It has been proposed that following light exposure, light induced hydrogen peroxide production in neurons will inactivate activated LITE-1, whose recovery then requires the thioredoxin and glutathione antioxidant systems (Fig. 1A and 1B) [12, 20-22]. Behavioural habituation to repeated light exposure is suppressed by the neuropeptide NLP-10 [17].

**Figure 1:**
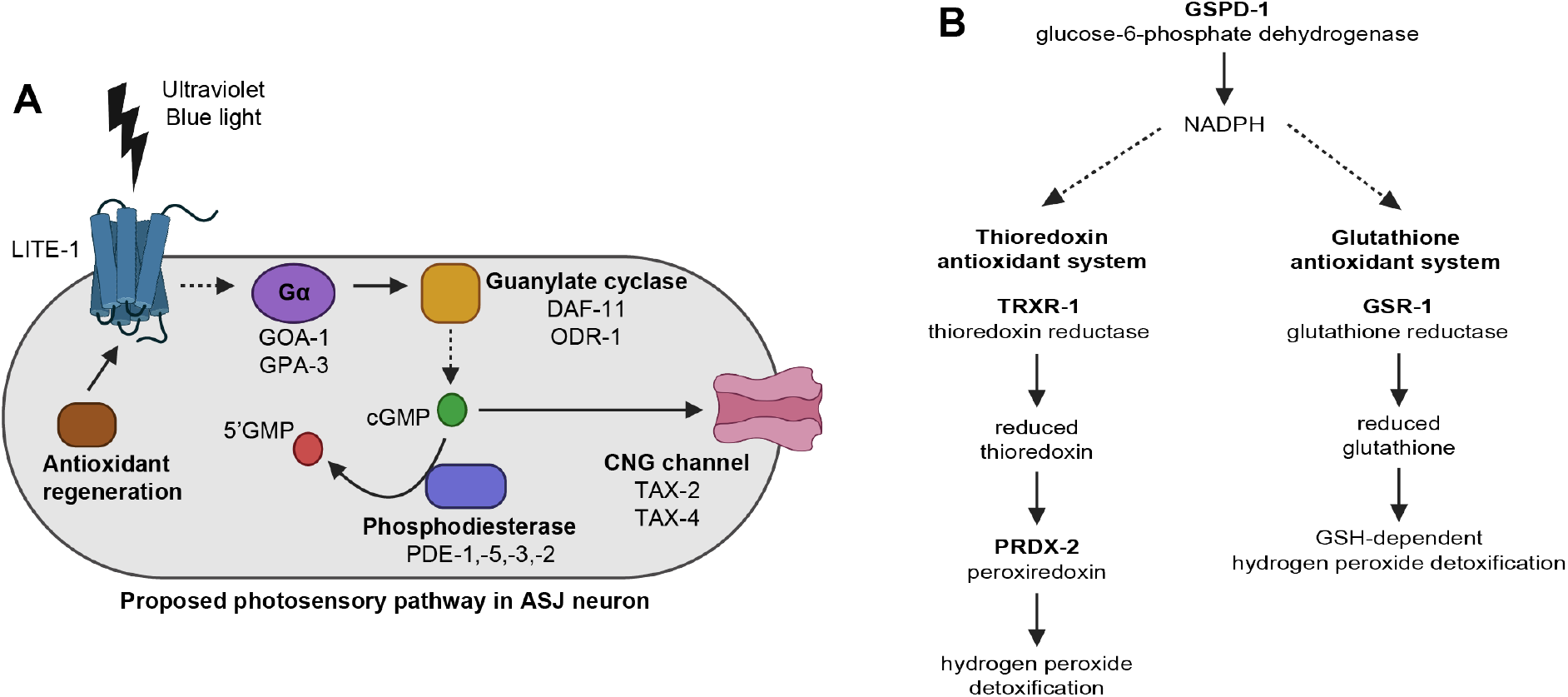
Proposed LITE-1-dependent photosensory response in *C. elegans* ASJ neuron. (A) Schematic of the proposed phototransduction signalling pathway mediating ultraviolet and blue light avoidance in *C. elegans* ASJ neuron. (B) Schematic of the thioredoxin and glutathione antioxidant systems. GSPD-1 generates NADPH required for both antioxidant systems. In the thioredoxin system, TRXR-1 regenerates reduced thioredoxin, which supports PRDX-2-dependent hydrogen peroxide detoxification. In the glutathione system, GSR-1 regenerates reduced glutathione, which supports GSH-dependent hydrogen peroxide detoxification [42-44]. Genes involved in the antioxidant systems tested in this study are highlighted in bold.

Here, we demonstrate that LITE-1-dependent photosensory function and avoidance of high concentration of diacetyl have partly different genetic requirements. Although phototransduction requires components of the cGMP signalling pathway, most of these were dispensable for diacetyl avoidance under the conditions tested.

## Results

### *goa-1* mutants show impaired avoidance of high concentration of diacetyl

LITE-1 mediates both light and high concentration diacetyl avoidance through ASK and ADL neurons, raising the question of whether these responses use shared or distinct downstream signalling components [9, 13, 15, 20]. To investigate this, we performed chemotaxis assays on the diacetyl avoidance response in worms with putative loss-of-function mutations in G proteins that were previously implicated in phototransduction, or expressed in at least one of the sensory neurons, ASK, ADL or ASH (Fig. 1A, 2A, S1A and S1B) [13, 23]. Among the candidates tested, mutants with loss-of-function in the *goa-1* gene exhibited a diacetyl avoidance defect comparable to that of the *lite-1* mutant (Fig. 2A). Multiple independent loss-of-function alleles of *goa-1* confirmed this phenotype (Fig. 2A).

**Figure 2:**
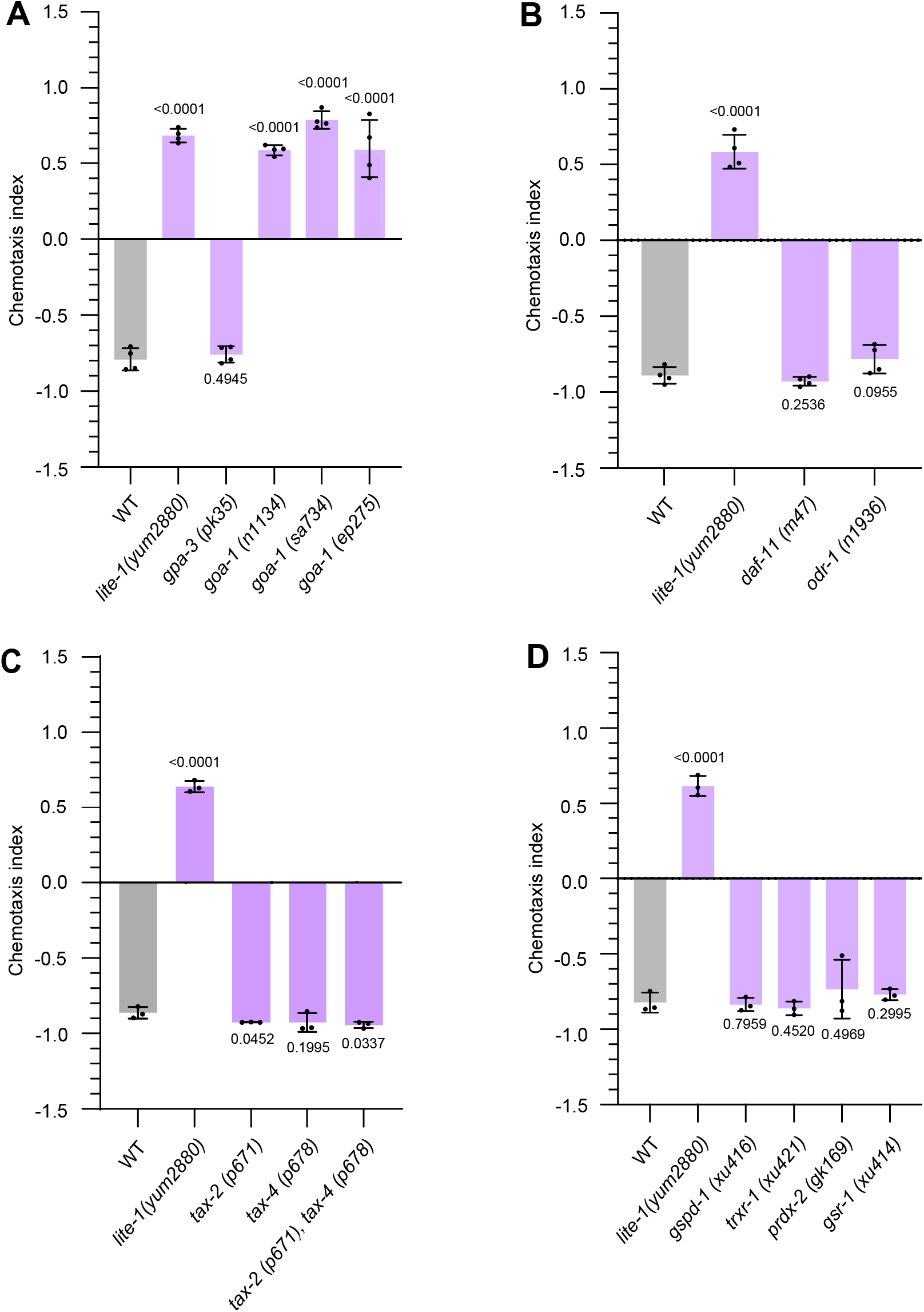
Diacetyl avoidance is retained in several photosensory signalling mutants. Chemotaxis index of wild-type animals and mutants affecting (A) G proteins, (B) guanylate cyclases, (C) CNG channels and (D) antioxidant systems in their response to undiluted diacetyl (5 μl). n = 3 or 4 biological replicates. Corresponding p-values are from unpaired two-tailed t-tests.

We next examined the loss of other core components that were previously implicated in photosensation such as guanylate cyclases (*daf-11* and *odr-1*), CNG channels (*tax-2* and *tax-*4), and the thioredoxin and glutathione antioxidant systems (Fig. 1A and 1B) [13, 20]. None of these mutants showed defects in the diacetyl avoidance behaviour (Fig. 2B, 2C and 2D). These results suggest that several genes required for normal photosensory responses are dispensable for the diacetyl avoidance behaviour.

### Altered adenylyl cyclase activity is associated with impaired diacetyl avoidance

GOA-1 can influence several intracellular signalling pathways, including pathways that regulate cyclic nucleotide levels [24, 25]. In *C. elegans*, ACY-1 is an adenylyl cyclase involved in cAMP signalling and functions downstream of GαS signalling [26, 27]. We thus tested a loss-of-function allele of *acy-1* which exhibited normal diacetyl avoidance, whereas two putative gain-of-function alleles displayed avoidance defects like those observed in *lite-1* mutants (Fig. 3A) [27, 28]. Supporting an association between adenylyl cyclase activity and diacetyl avoidance, treating wild-type animals with forskolin, an activator of adenylyl cyclase, also impaired diacetyl avoidance (Fig. 3B) [28-30].

**Figure 3:**
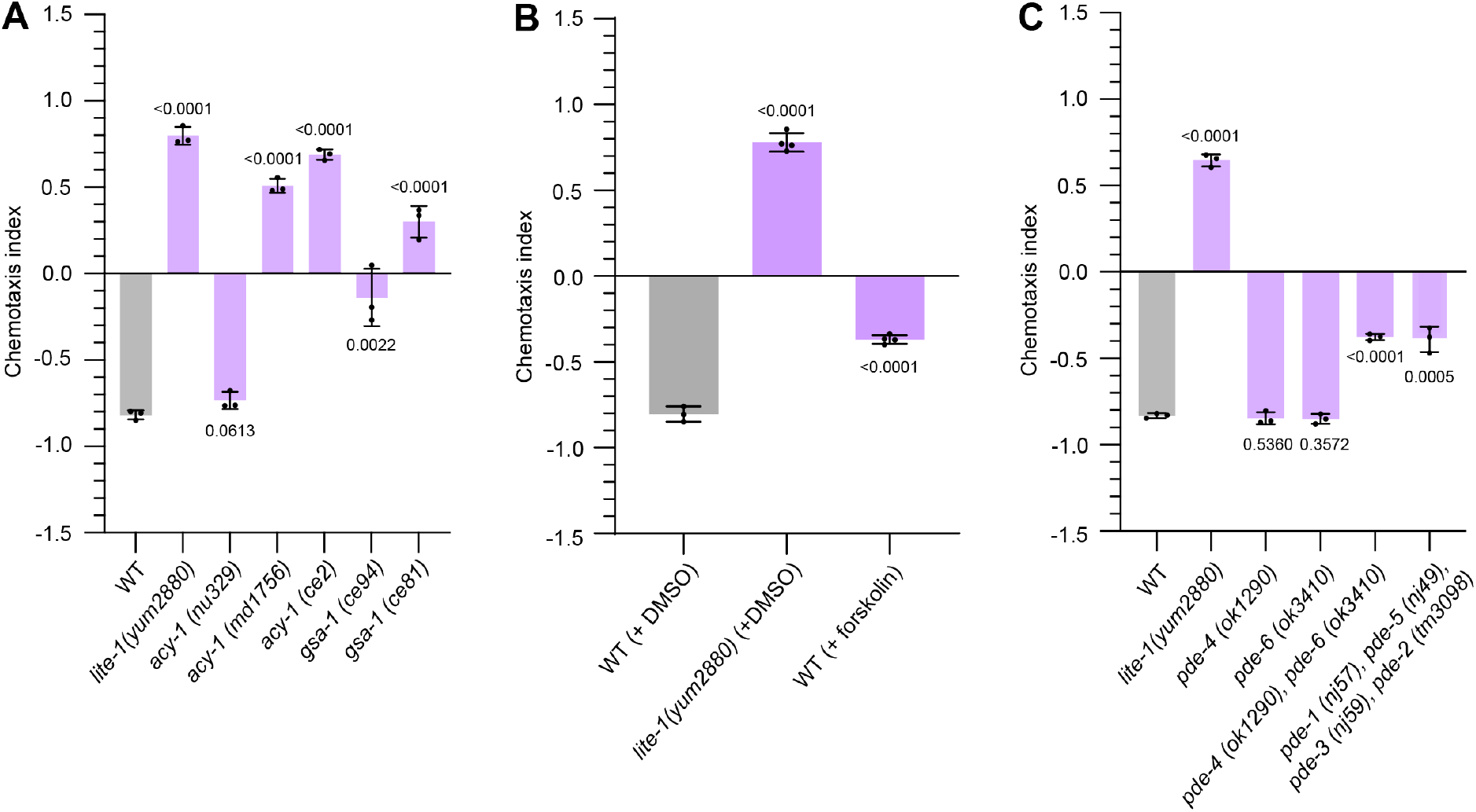
Altered adenylyl cyclase activity is associated with impaired diacetyl avoidance. Chemotaxis index of wild-type animals and mutants affecting (A) adenylyl cyclase and GSA-1 signalling, (B) wild-type animals treated with 80 μm forskolin or 0.16% DMSO as vehicle control and, (C) phosphodiesterases in their response to undiluted diacetyl (5 μl). n = 3 biological replicates. Corresponding p-values are from unpaired two-tailed t-tests.

To further examine the association between altered adenylyl cyclase activity and the avoidance response, we next analysed two gain-of-function alleles of *gsa-1*, which encodes GαS, a stimulator of ACY-1 activity [25, 27]. Both mutants of *gsa-1* showed significant defects in diacetyl avoidance (Fig. 3A).

We next asked whether disrupting cyclic nucleotide regulation more broadly also affects the avoidance behaviour. To address this, we examined mutants in phosphodiesterases (PDEs), which regulate cAMP and/or cGMP levels [28, 31, 32]. Disruption of several PDE genes resulted in diacetyl avoidance defects (Fig. 3C). These phenotypes suggest that altered cyclic nucleotide regulation can influence the avoidance behaviour, although it should be noted that individual PDEs differ in their substrate preferences and have broad cellular functions.

### Cilia defective mutants retain normal diacetyl avoidance behaviour

LITE-1-dependent avoidance of high concentration of diacetyl is mediated by the ASK and ADL neurons, both of which possess ciliated sensory endings [9]. Because defective cilia frequently impair chemosensory behaviours, we next examined whether disruptions of cilia affect the diacetyl avoidance behaviour. We screened a panel of cilia defective mutants with disruption in components of the intraflagellar transport (IFT) machinery, which is essential for the assembly and maintenance of cilia [33, 34]. The mutants analysed include *osm-3* (kinesin motor), *klp-11* (kinesin-II motor), *kap-1* (kinesin-associated protein), *che-3* (dynein motor), *daf-10* (IFT-A complex), *osm-5* and *osm-6* (IFT-B complex). None of the tested mutants exhibited significant defects in diacetyl avoidance, indicating that intact cilia are not strictly required for the avoidance behaviour, as has also been reported for LITE-1-dependent photosensation (Fig. 4A) [18].

**Figure 4:**
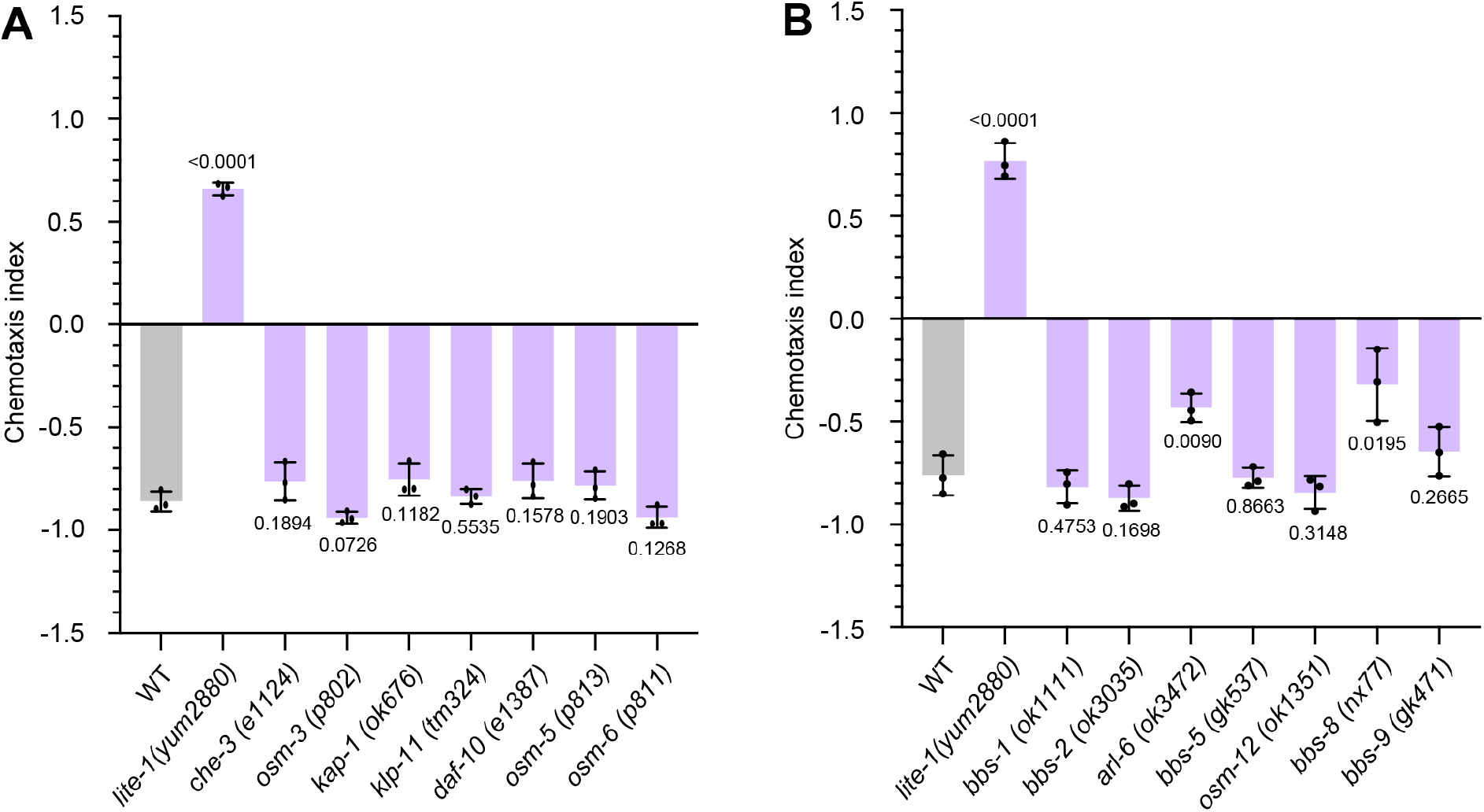
Cilia defective and BBSome-related mutants retain diacetyl avoidance. Chemotaxis index of wild-type animals and mutants in (A) intraflagellar transport machinery and (B) BBSome complex in their response to undiluted diacetyl (5 μl). OSM-12 is also known as BBS-7 [45]. n = 3 biological replicates. Corresponding p-values are from unpaired two-tailed t-tests.

Although disruption of the IFT machinery did not affect diacetyl avoidance, other cilia associated proteins may regulate LITE-1 function, such as the BBSome which has been implicated in the regulation of LITE-1 photosensation [18]. We thus examined BBS-related mutants, of which *arl-6* and *bbs-8* mutants exhibited partial defects in diacetyl avoidance (Fig. 4B). However, because these mutants also display locomotion defects, it remains unclear whether the observed phenotypes reflect altered sensory function or impaired movement.

Nevertheless, these findings indicate that most of the mutants affecting cilia structure or BBSome function retain normal avoidance of high concentration of diacetyl, suggesting that intact cilia and canonical BBSome function are not broadly required for the diacetyl response.

## Discussion

The LITE-1-mediated photosensory response involves a well characterised cGMP-dependent signalling cascade that comprises G proteins, guanylate cyclases, CNG channels, phosphodiesterases and antioxidant systems that drive the light evoked avoidance response (Fig. 1A and 1B) [13, 15, 20]. By contrast, our behavioural analyses revealed that several components required for LITE-1 photosensation were dispensable for avoidance of high concentration of diacetyl. These findings suggest that LITE-1-dependent photosensation and diacetyl avoidance have at least partly different genetic requirements.

It is possible that this divergence from phototransduction pathway may reflect a functional adaptation that allows LITE-1 to respond to two fundamentally different environmental cues through distinct signalling mechanisms. The use of distinct signalling mechanisms for different stimuli is consistent with observations from other sensory receptors. The sweet taste receptor heterodimer T1R2/T1R3 responds to a broad range of sugars and sweeteners by producing distinct intracellular signalling profiles depending on the ligands it encounters [35]. Sucralose has been reported to induce both Ca^2+^and cAMP responses, glycyrrhizin predominantly promotes Ca^2+^signalling, whereas saccharin increases cAMP levels [35]. These ligand dependent outputs then engage different downstream pathways, including PLCβ2/IP_3_ signalling and cAMP/PKA signalling [35, 36].

Whether LITE-1 employs a similar strategy to sense and distinguish different sensory inputs remains to be determined. Future studies using *lite-1* mutants targeting specific functional domains, such as channel gating, ligand binding pocket, or light sensitive residues, may clarify how LITE-1 contributes to both responses [12, 14]. Tryptophan residues not required for light sensing (W41, W191, W319, and W419) are possible targets to assess their essentiality for diacetyl detection [12, 14]. However, such experiments would need to establish whether the expression levels and localisation of the mutant LITE-1 remain unchanged, so as to distinguish functional effects from indirect effects such as altered protein stability. Additionally, differences between the two avoidance behaviours could also arise at the circuit level, where recent work has shown that the LITE-1-expressing interneuron AVG responds to blue light and releases the neuropeptide NLP-10, which acts to suppress behavioural habituation to repeated light exposure [17]. Whether NLP-10 is also involved in responses to sustained diacetyl exposure remains to be tested.

The data we have presented here can suggest hypotheses but do not support firm conclusions without further work because of several limitations. First, many of the examined genes have broad roles in neuronal signalling and synaptic transmission. We, therefore, could not exclude the possibility that some of the observed mutant phenotypes are the result of general sensory impairment. The testing of diacetyl defective mutants to a range of attractant or repellent odorants will be necessary to address this possibility. Second, parallel sensory pathways may mask the contribution of individual signalling genes to LITE-1-dependent responses. We have recently found that avoidance of high concentration of diacetyl may require both ASK and ADL neurons [9]. The avoidance defect may arise through as yet unidentified receptors or putative low-affinity diacetyl receptors, or through shared downstream sensory circuitry [8]. Third, the current study relies heavily on endpoint behavioural assays. Future studies that directly measure neuronal activity in the avoidance defective mutants will be informative in determining how these genes are involved in the response. Fourth, a genetic screen for mutant worms that fail to avoid high concentration of diacetyl but still retain normal response to light will be a powerful approach to identify components involved in LITE-1-dependent chemosensation. Finally, the defects we have identified in the tested strains cannot be confidently attributed to the mutated genes without rescue experiments.

Despite these limitations, on balance, our findings suggest that avoidance of high concentration of diacetyl has distinct genetic requirements from LITE-1-dependent photosensation. While GOA-1 and cyclic nucleotide regulation are likely important for both light and diacetyl avoidance, most of the canonical components of the phototransduction pathway and intact sensory cilia are largely dispensable for diacetyl avoidance. These results suggest the hypothesis that LITE-1 may engage different downstream mechanisms for different stimuli, a possibility that remains to be tested.

## Methods

### Worm strains and bacterial maintenance

The *E. coli* wild-type isolate OP50 was obtained from Caenorhabditis Genetics Center (CGC) and maintained under standard laboratory conditions on lysogeny broth agar. Worm strains were maintained as previously described at 20°C on nematode growth medium seeded with *E. coli* wild-type isolate OP50 [37]. The worm strains used in this study were obtained from the CGC, unless otherwise stated, N2; wild-type isolate, CHS1610; *lite-1(yum2880) X* [38], NL335; *gpa-3(pk35) V*, TM2426; *goa-1(n1134) I*, DG1856; *goa-1(sa734) I*, CE1051; *goa-1(ep275) I*, DR47; *daf-11(m47) V*, CX2065; *odr-1(n1936) X*, PR671; *tax-2(p671) I*, PR678; *tax-4(p678) III*, BR5514; *tax-2(p671) I; tax-4(p678) III*, TQ10548; *trxr-1(xu421) IV*, VC289; *prdx-2(gk169) II*, TQ10521; *gsr-1(xu414) III*, TQ10517; *gspd-1(xu416) IV*, RB1231; *pde-4(ok1290) II*, RB2470; *pde-6(ok3410) I*, CHS785; *pde-4(ok1290) II; pde-6(ok3410) I* [39], TQ1828; *pde-1(nj57) I; pde-5(nj49) I; pde-3(nj59) II; pde-2(tm3098) III*, KP1182; *acy-1(nu329) III*, KG522; *acy-1(md1756) III*, KG518; *acy-1(ce2) III*, KG524; *gsa-1(ce94) I*, KG421; *gsa-1(ce81) I*, CB1124; *che-3(e1124) I*, PR802; *osm-3(p802) IV*, RB849; *kap-1(ok676) II*, VC1228; *klp-11(tm324) IV*, CB1387; *daf-10(e1387) IV*, PR813; *osm-5(p813) X*, PR811; *osm-6(p811) V*, VC837; *bbs-1(ok1111) I*, RB2242; *bbs-2(ok3035) IV*, RB2509; *arl-6(ok3472) III*, VC1316; *bbs-5(gk537) III*, RB1268; *osm-12(ok1351) III*, MX52; *bbs-8(nx77) V*, VC1062; *bbs-9(gk471) I*, CX3222; *odr-3(n1605) V*, NL332; *gpa-1(pk15) V*, NL1137; *gpa-5(pk376) X*, NL1146; *gpa-6(pk480) X*, NL787; *gpa-11(pk349) II*, NL2330; *gpa-13(pk1270) V*, NL788; *gpa-14(pk347) I*, NL797; *gpa-15(pk477) I*, NL792; *gpc-1(pk298) X*.

### Chemotaxis assay

Chemotaxis assays were performed under ambient light, with care taken to avoid exposure to any strong light source. The assays were performed in 9 cm petri dishes filled with 10 ml of agar (1 mM MgSO_4_, 1 mM CaCl_2_, 1.7% agar and 25 mM K_2_HPO_4_ (adjusted to pH 6 with KOH)) at 20°C as previously described [40]. In brief, for the plate layout, two parallel lines with 1 cm apart were drawn onto the middle where the animals will be placed. On each opposite side of the plate, two spots were designated for odorant and control (Fig. S1A). 1 μl of 1 M NaN_3_ (Sigma-Aldrich, catalogue number: 71290) was then added onto the four spots to immobilise the animals. 5 μl of diacetyl (Thermo Fisher Scientific, catalogue number: 107651000) and 5 μl of ethanol as control (Thermo Fisher Scientific, catalogue number: 16688242) were then spotted onto the respective dots. Synchronized animals were obtained as previously described [41], with L1 larvae in diapause for 48 h before refed onto NGM plates seeded with *E. coli* OP50 and maintained at 20°C. Synchronized day one adults were then washed 3 times in chemotaxis buffer (1 mM MgSO_4_, 1 mM CaCl_2_ and 25 mM K_2_HPO_4_ (adjusted to pH 6 with KOH)) before being transferred onto the centre of the plate and the chemotaxis assay will last for 1 hr at 20 °C. The chemotaxis plates were then stored at 4 °C before counting. The chemotaxis index was calculated as (number of animals on the odorant side – number of animals on the control side) / (sum of all animals on both sides) (Fig. S1A). Animals between the parallel lines were excluded. A chemotaxis value of +1 indicates that all animals were within the odorant side (N^+^), while a chemotaxis value of -1 indicates that all animals were within the control side (N^-^). Care was taken to minimise the exposure of the diacetyl stock to light.

### Forskolin treatment

Forskolin was obtained from Sigma-Aldrich (catalogue number: 344282) as a 50 mM stock solution in DMSO, and treatment of animals were as previously described [30]. Synchronized day one adults were washed 3 times in M9 medium before treatment with 80 μm forskolin (final concentration of DMSO at 0.16%) and OP50 in M9 medium for 2 hrs at 20°C on a rotator. For vehicle control, animals were treated with 0.16% DMSO under the same condition. After treatment, animals were transferred directly onto chemotaxis plates, and chemotaxis assays were performed as described above.

### Statistical analysis

Each chemotaxis experiment was performed with 3 to 4 biological replicates conducted on different days. Each dot on the graph represents a biological replicate, calculated as the mean of 4 plates. Mutant animals were analysed on the same day with their corresponding control animals. Statistical analyses were performed with GraphPad Prism 11 using unpaired two-tailed t-tests, with statistical significance shown as p-values in their corresponding graphs.

## Acknowledgements

We thank Na Yu for technical assistance. Flint Stevenson-Jones and Hippo family for support. This work was funded by the Medical Research Council (grant MC-A658-5TY30 to AEXB). Some strains were provided by the CGC which is funded by the NIH Office of Research Infrastructure Programs (P40 OD010440).

## Author contributions

A.K.: Conceptualization, data curation, formal analysis, investigation, validation, methodology, visualization, writing – original draft and writing – review and editing. C.C.: Conceptualization, methodology and writing – review and editing. A.E.X.B.: Supervision, project administration, conceptualization, methodology, writing – review and editing, funding acquisition.

## Data availability statement

The datasets generated in this publication are available upon request.

---

**Figure S1:**
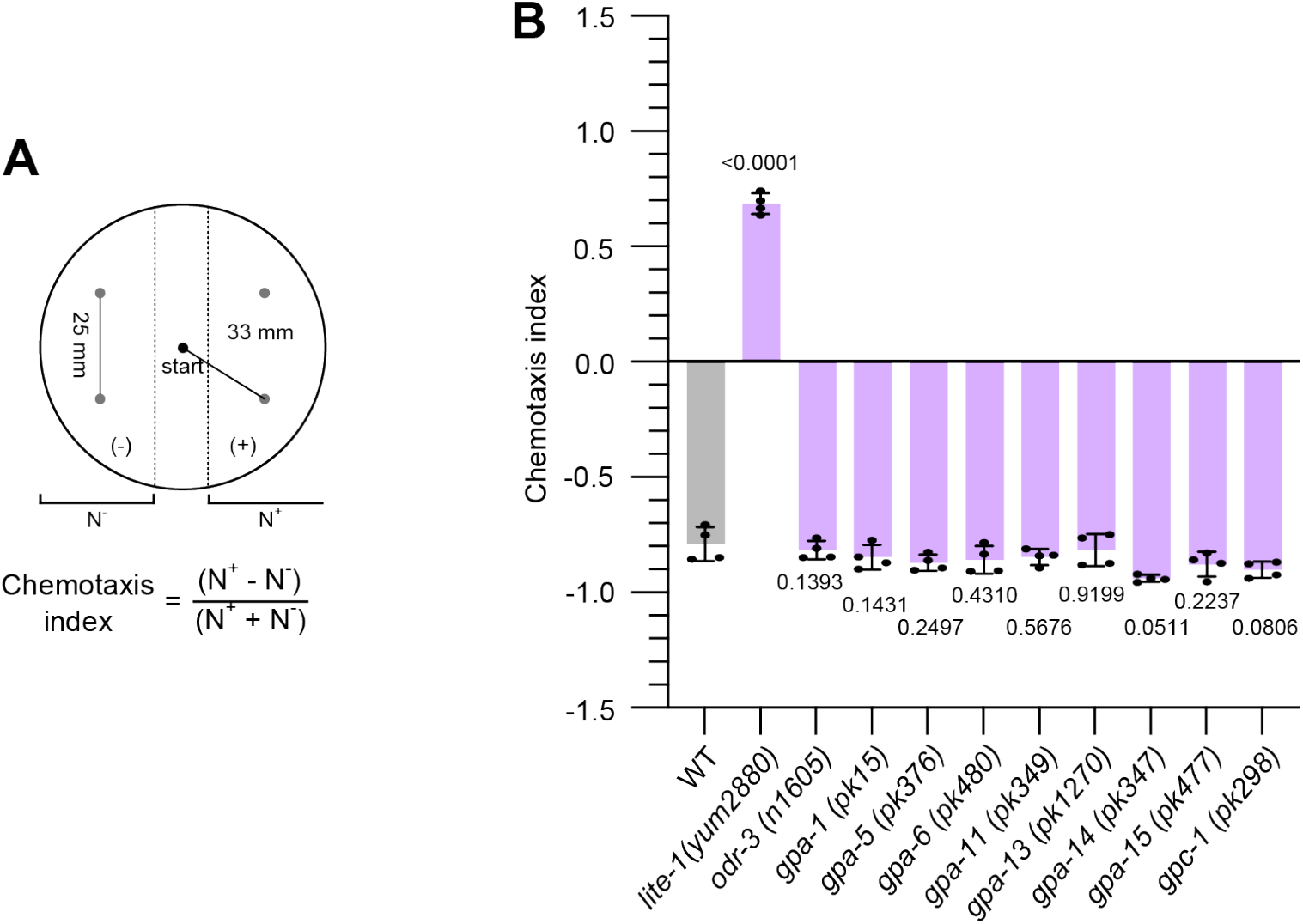
Diacetyl avoidance in additional G protein mutants. (A) Schematic of chemotaxis assay setup as previously described [40]. N^+^ and N^-^ indicate the odorant and control side respectively. (B) Chemotaxis index of wild-type animals and G protein mutants in their response to undiluted diacetyl (5 μl). n = 4 biological replicates. Corresponding p-values are from unpaired two-tailed t-tests.

